# Innate immune signalling drives loser cell elimination during stem cell competition in the *Drosophila* testis

**DOI:** 10.1101/2020.03.05.979161

**Authors:** Silvana Hof-Michel, Ljubinka Cigoja, Sabina Huhn, Christian Bökel

## Abstract

In the *Drosophila* testis, a group of stromal cells termed hub provides multiple niche signals for the surrounding germline and somatic stem cells. Stem cells of both populations compete for physical retention in the niche, and stem cell clones unable to transduce any one niche signal are rapidly eliminated by differentiation. We have recently mapped the transcriptomes of isolated somatic cyst stem cells and differentiated cyst cells, and found that the stem cells but not their differentiated progeny activate an immune response involving the NF-κB transcription factor Relish (Rel).

Here we show i) that Rel activation is not required for stemness but occurs physiologically in “losers” of stem cell competition, ii) that loss of Rel or the upstream receptor Toll suppresses loser elimination irrespective of how loser fate was induced, and iii) that clonal Rel activation is sufficient for the displacent of neutral or winner cells from the niche, even if the winners otherwise retain stem cell properties.

This generalized mechanism for the elimination of “loser” stem cells may mask the compound nature of stem cell behaviour, and instead generate the impression of a binary cell fate decision between stemness and differentiation.

## Introduction

Cell competition is a process of tissue quality control whereby adjacent cells compare their relative fitness, resulting in the active elimination of viable but less fit “loser” cells from the tissue by the more fit “winner” cells (Bowling et al., 2019; Madan et al., 2018). Cell competition in this narrow sense is distinct from neutral competition, where cells achieve clonal dominance by - potentially only transiently or stochastically - exhibiting a higher proliferation rate than their otherwise equally fit neighbours (de Navascues et al., 2012; Snippert et al., 2010). While cell competition has been implicated in processes like tissue homeostasis, delay of tissue aging, and both cancer prevention and oncogenesis in a wide range of organisms, many of the underlying principles have initially been uncovered using *Drosophila* as a model system (Bowling et al., 2019; Madan et al., 2018). In particular, work in larval imaginal discs has identified various parameters that contribute to the overall cellular fitness that is measured during competition, including differences in the capability to transduce growth or survival signals (Moreno et al., 2002), heterozygosity for mutations in ribosomal proteins (Minutes) (Simpson and Morata, 1981), supercompetition by cells with elevated myc expression (de la Cova et al., 2004; Moreno and Basler, 2004), or loss of epithelial integrity (Tamori et al., 2010).

Genetic experiments in the imaginal discs already demonstrated that various components of the Toll and Imd pathways, the two main signalling cascades of the *Drosophila* innate immune response (Ferrandon et al., 2007), are also involved in cell competition (Meyer et al., 2014). In the immune context, the Imd pathway transduces signals from surface peptidoglycan receptors through the intracellular death domain adaptor Imd, ultimately activating the NF-κB family transcription factor Rel by phosphorylation and proteolytic removal of its C-terminal autoinhibitory domain (Wiklund et al., 2009). In contrast, the canonical Toll (Tl) pathway is, in the immune context, triggered by proteolytic activation of the ligand Spätzle (Spz) by protease cassettes acting as pathogen sensors. Activated Spz induces clustering of the transmembrane receptor Tl and subsequently the phosphorylation and degradation of the IκB homologue Cactus that retains the NF-κB transcription factors Dl or Dif in the cytoplasm. Once in the nucleus Dl, Dif, or the active, N-terminal fragment of Rel regulate pathway specific expression of antimicrobial peptides and other target genes (Ferrandon et al., 2007).

In the competition context, separation between the two pathways seems to be less clear cut, with components instead assembled into “competitive signalling modules” (Alpar et al., 2018) whose composition depends on the specific mechanism triggering competition: In the wing disc, competition between Minute (RpL14^+/−^) loser cells and their wild type neighbours e.g. involves a module comprising Spz, Tl, and the related receptors Toll-3 and Toll-9 that activates Dl and Dif in the losers and triggers their apoptosis through transcription of Rpr (de la Cova et al., 2014; Meyer et al., 2014). In contrast, elimination of WT wing disc cells by Myc overexpressing supercompetitor neighbours relies on the atypical activation of the canonical IMD target Rel through a cassette containing Tl, Toll-2, −3, −8, and −9, which then drives expression of the pro-apoptotic factor Hid in the wild type losers (Meyer et al., 2014).

Cell competition has also been observed in the stem cell niche of the fly testis. There, germline stem cells and somatic cyst stem cells surround a cluster of postmitotic somatic cells termed hub that provides the niche signals for the maintenance and activity of both stem cell populations. Access to this niche is a limited resource, hence stem cells compete both between and within the two stem cell pools for contact with the hub. Intriguingly, the result of competition between these stem cells is generally not apoptosis of the losers, but their elimination from the stem cell compartment by differentiation (Fig. 1A).

**Figure 1:**
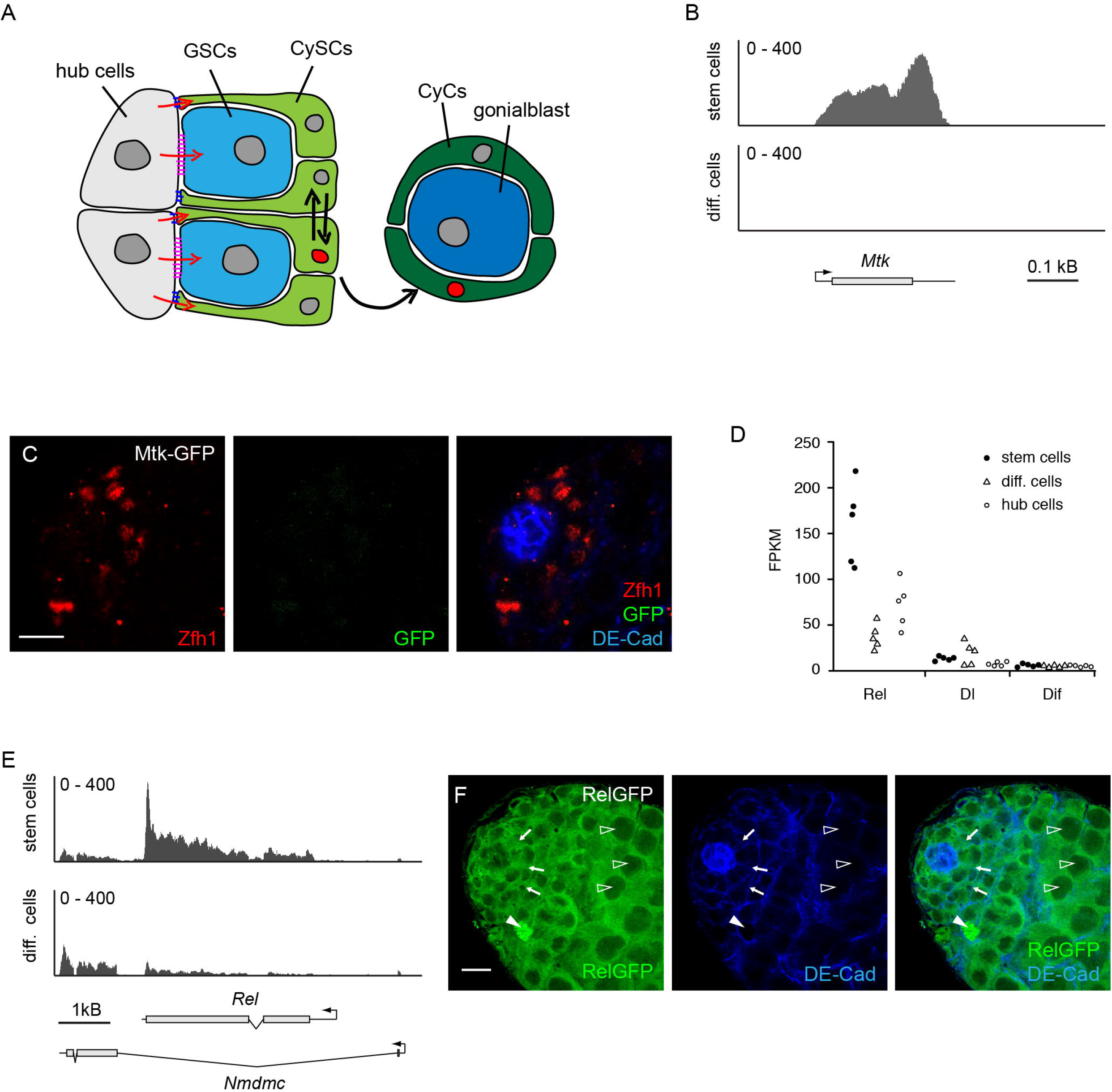
Expression of innate immune signalling genes in the *Drosophila* testis. (A) Schematic representation of the testis tip. The hub (grey) provides niche signals (red arrows) to germline stem cells (GSCs, light blue) and somatic cyst stem cells (CySCs, light green) with which it stays in contact through adhesion junctions (red and blue) and which give rise to gonialblast (dark blue) and their ensheathing cyst cells (CyCs, dark green), respectively. Somatic cells compete for retention in the stem cell pool, and loser cells (red nucleus) are preferentially forced to differentiate (black arrows). (B) RNAseq reveals expression of the antimicrobial peptide Mtk in isolated CySCs (top track) but not differentiated CyCs (bottom track). (C) In testes fixed immediately after dissection a Mtk-GFP reporter (green) is not expressed in the vicinity of the hub (marked by DE-Cad, blue) or the Zfh1 positive (red) pool comprised of CySCs and their immediate, differentiated cyst cells progeny. (D) Expression levels in FPKM for the three *Drosophila* NF-κB transcription factors in the somatic lineage of the testis. Each data point represents one replica transcriptome. (E) Rel expression is upregulated in CySCs (top track) relative to differentiated CyCs (bottom track). (F) A GFP-tagged BAC construct reveals Rel expression in CySCs (arrows) as well as the germline. With the exception of a few dying cells (solid arrowhead), Rel-GFP is excluded from the germline nuclei (open arowheads). Hub and cell outlines marked by DE-Cad (blue). (B,E) Number of reads plotted against genome position, (C, F) Scale bars 10 μm.

The endogenous equilibrium, where occupancy of the hub surface is strongly biased towards the GSCs, can be modified by manipulating the Suppressor of Cytokine Receptor Signalling 36 (SOCS36) (Issigonis et al., 2009). Even though adhesion and niche occupancy by somatic cyst stem cells were therefore initially linked to Jak/Stat activity, more recent results suggest that the relative competitive strength of individual somatic stem cells is instead set by MAPK signalling (Amoyel et al., 2016; Singh et al., 2016). In addition, competitive ability of somatic cyst stem cells depends on Cadherin mediated adhesion to the hub that is regulated through a Slit/Robo2/Abl axis (Stine et al., 2014).

Similar to the wing disc epithelium, where cells compete for survival factors like Dpp (Moreno et al., 2002), differences in niche signalling activity are key parameters defining the relative fitness of individual stem cells in the testis. In the somatic lineage, the Hh (Amoyel et al., 2013; Michel et al., 2012) and Upd - Jak/Stat (Kiger et al., 2001; Tulina and Matunis, 2001) niche signalling cascades act in parallel. While both pathways have unique targets they also converge on the key stem cell transcription factor Zfh1 (Leatherman and Dinardo, 2008). Loss of Zfh1 or inactivation of either of the niche signalling pathways causes the affected cells to lose stem cell status. In case of Hh in particular it is unambiguous that this elimination process involves cell competition: Individual cells in which the ability to transduce the Hh signal is clonally inhibited, e.g. by mutation of the key signal transducer Smoothened (Smo), are rapidly eliminated by differentiation (Michel et al., 2012). In contrast, global inactivation of Hh signalling using a temperature sensitive Hh allele has only minor effects, and stem cells are maintained long term in the absence of Hh activity (Michel et al., 2012). Conversely, clonal activation of the Hh signal causes the affected stem cells to outcompete their heterozygous sibling cells from the niche by skewing the ongoing neutral competition (Amoyel et al., 2013; Amoyel et al., 2014; Michel et al., 2012).

Participation in competition with neighbouring stem cells, including specifically the propensity to be forced into differentiation when found in a loser state, is thus a hallmark of gonadal stem cell physiology, comparable to other aspects of stemness such as e.g. adhesion to the niche, proliferation, absence of differentiation, or a stem cell specific metabolic mode.

As reported in detail in a separate manuscript (Hof-Michel and Bökel, 2020), we have mapped the transcriptomes of isolated somatic cyst stem cells and their differentiated cyst cell progeny by RNAseq. Intriguingly, genes differentially upregulated in stem cells were highly enriched for genes involved in innate immunity, including signalling pathway components as well as antimicrobial peptide effectors, indicating an active immune response. It appears implausible that the stem cells, but not the immediately adjacent differentiated cyst cells should have a need for antimicrobial activity under physiological conditions. However, even if the observed antimicrobial activity were an artefact of the isolation procedure, the differential ability of the stem cells to trigger such an immune response must still be accounted for.

The gene encoding the NF-κB transcription factor Rel was also amongst the strongest hits in a previous screen for Zfh1 binding in the somatic cyst stem cells, marking Rel as a potential target of niche activity (Albert et al., 2018). We therefore examined a potential link between immune signalling involving Rel, niche function, and the previously observed stem cell competition.

Here we report that transient Rel activation occurs in cells adopting “loser” fate during stem cell competition. Loser cells with a compromised ability to transduce the Hh or Upd-Jak/Stat niche signals or lacking certain stem cell expressed transcription factors that are normally outcompeted are retained in the stem cell population following concomitant inactivation of Rel. Clonal loss of Tl, but not of the canonical Rel upstream signal transducer IMD, elicits the same effect. Rel is thus required in the losers for their elimination, and its activation appears to involve a “competitive signalling module” (Alpar et al., 2018) downstream of Tl.

Expression of a constitutively active form of Rel is sufficient to trigger the loss of neutral clones from the niche. It is also able to induce displacement of winner cells from the niche, without, however, affecting the increased winner clone size or the expression of stem cell markers. Constitutive Rel activation thus uncouples physical retention of stem cells within the niche from other aspects of stemness.

We therefore propose i) that the cell type specific expression of Rel and other components of the innate immune cascade renders cyst stem cells sensitive to competitive elimination from the niche and differentiation, ii) that participation in this quality control mechanism presents a further, genetically separable aspect of stemness, and iii) that this shared mechanism for the elimination of loser cells, irrespective of how their fitness was compromised, may mask the compound nature of stemness and generate the impression of a binary decision between stem cell fate and differentiation.

## Results

### Isolated CySCS but not their differentiated progeny are able to activate an innate immune response

As described in detail in an accompanying manuscript (Hof-Michel and Bökel, 2020), RNAseq of purified somatic *Drosophila* testis cells demonstrated that the transcriptome of CySCs is, relative to that of the differentiated CyCs, highly enriched in genes involved in immune / NF-κB signalling and antimicrobial activity (suppl. Table S1). Stem cells but not their differentiated progeny e.g. express the antimicrobial peptides Metchnikovin (Mtk) (Fig. 1B) or Drosomycin (Drs) (Fig. S 1A). However, cell sorting by FACS required mechanic and enzymatic isolation of the cells of interest. Exposure to bacteria, in particular from the ruptured intestine, prior to RNA extraction was therefore unavoidable.

To check whether cyst stem cells activated immune signalling also under physiological conditions we imaged testes from established GFP reporter lines for Mtk (Clark et al., 2013) and Drs (Ferrandon et al., 1998). However, neither line showed any GFP signal in the vicinity of the niche when the testes were fixed immediately after dissection (Fig. 1C, Fig. S1B). This negative result was not unexpected, since it appeared implausible that the cyst stem cells should, under physiological conditions, have a need to respond to pathogens, while their immediate progeny, with which they remain in close contact, do not. Nevertheless, the striking observation that the two related, somatic cell populations differ in their ability to activate the relevant upstream pathways when exposed to pathogens during isolation remains and requires an explanation.

### Loss of Rel suppresses the elimination of loser cells with compromised Hh niche signal transduction

Considering that the somatic stem cells of the testis niche experience strong competition (Amoyel et al., 2016; Amoyel et al., 2013; Amoyel et al., 2014; Issigonis et al., 2009; Michel et al., 2012; Stine et al., 2014) and that the Tl and Imd pathways are involved in cell competition in epithelia (Alpar et al., 2018; Meyer et al., 2014), we instead tested a possible requirement for innate immune signaling in stem cell competition. We focussed on Rel, as it was the most highly expressed of the three Drosophila NF-κB transcription factors in the somatic lineage of the testis and, in particular, the cyst stem cells (Fig. 1D,E). However, in intact testes the somatic expression of Rel was largely masked by Rel expression in the germline (Fig. 1F). We therefore instead directly turned to MARCM (Lee and Luo, 1999) analysis of Rel mutant somatic stem cells.

Clonal analysis of somatic stem cell behaviour in the adult testis is greatly facilitated by the fact that the stem cells are the only proliferating cells within this lineage: Any observed clone must therefore have been born in a stem cell division (Le Bras and Van Doren, 2006; Sinden et al., 2012). Homozygous *smo* MARCM clones created in an otherwise heterozygous background during CySC division adopt “loser” fate and are rapidly and quantitatively eliminated from the stem cell pool by differentiation (Michel et al., 2012). In contrast, inactivating Hh signalling in all cells using the temperature sensitive Hh allele *hh*^*ts2*^ only has a minor effect: Even though their numbers are somewhat reduced, somatic stem cells are maintained over extended periods in the absence of Hh signalling, and the flies remain fertile (Michel et al., 2012). The rapid elimination of *smo* mutant clones is therefore unambiguosly caused by cell competition.

However, *smo* is located on chromosome arm 2L, while *rel* and other innate immunity genes of interest are located elsewhere in the genome. This makes the testing of genetic interactions by MARCM using *smo* loss of function mutations experimentally inconvenient. We therefore turned to clonal overexpression of a non-phosphorylatable, dominant negative, and hence nonfluorescent Smo activation sensor (Smo-InversePericam-SA, here abbreviated as SmoDN) (Kupinski et al., 2013) to cell autonomously block Hh signalling and induce loser fate. Similar to globally abolishing Hh signalling with the help of *hh*^*ts2*^, overexpressing SmoDN in all cells of the cyst cell lineage under tj-Gal4 tub-Gal80^ts^ control only had a minor effect on the number of Zfh1 positive cells per testis (Fig. S2A). In contrast, and consistent with a competition scenario, the fraction of SmoDN overexpressing homozygous clonal cells retained in the Zfh1 positive compartment containing the stem cells and their most recent progeny was severely reduced relative to control by 5d after clone induction (ACI) (Fig. 2A-C). We could thus test the effect of Rel inactivation on cell competition by expressing SmoDN in MARCM clones for the loss of function allele *rel*^E20^ that eliminates expression of all Rel isoforms (Hedengren et al., 1999). Homozygous *rel*^E20^ flies are viable and fertile, demonstrating that Rel is not by itself needed for stem cell function. Consistently, *rel*^E20^ clones were retained in the stem cell compartment with similar frequency as control clones (Fig. 2D,F). Importantly, *rel*^E20^ clones overexpressing SmoDN retained a significantly larger fraction of Zfh1 positive cells than neutral control clones overexpressing SmoDN (Fig. 2E,F). Thus, loser cells with impaired ability to transduce the Hh niche signal require Rel for their effective elimination by differentiation.

**Figure 2:**
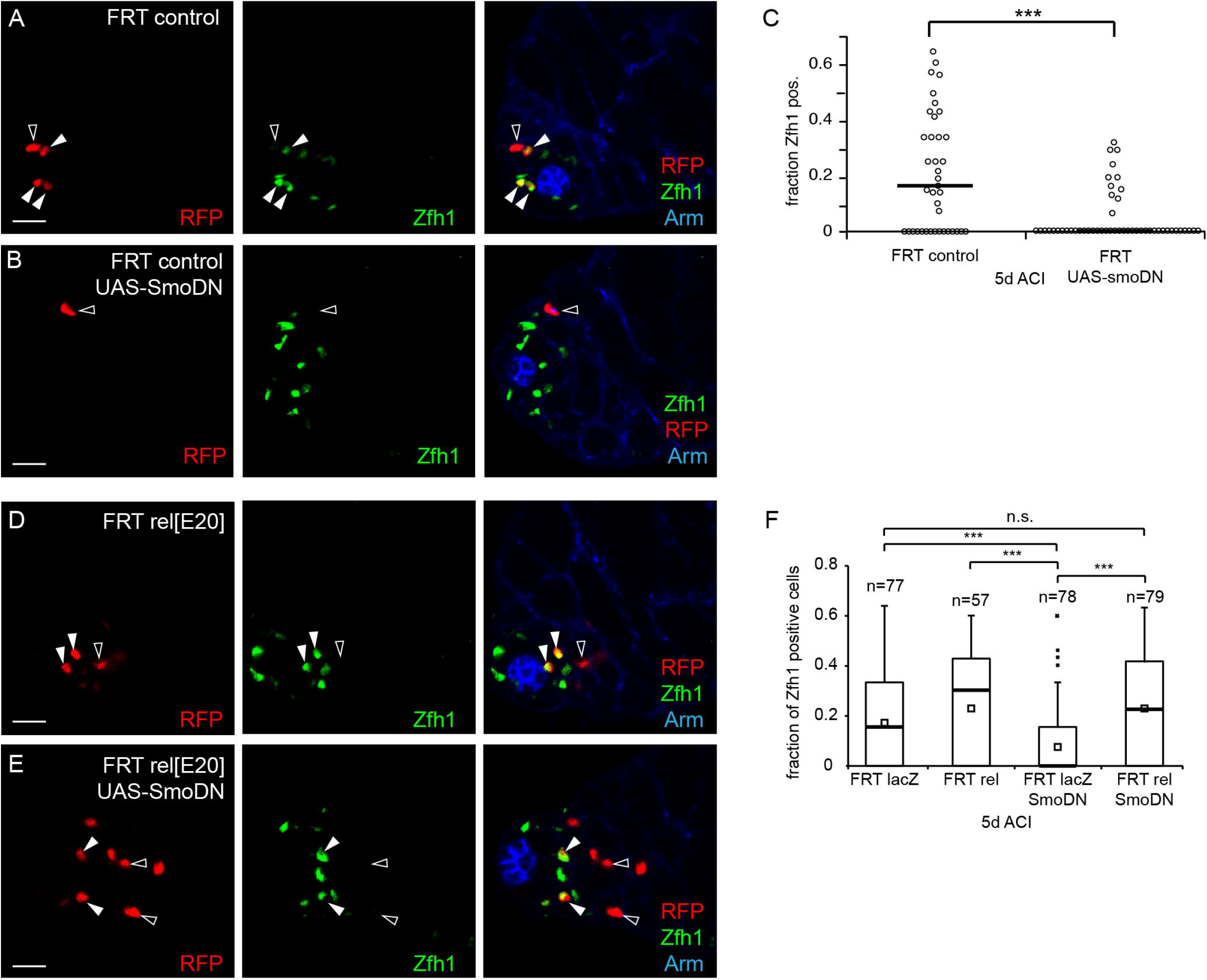
MARCM analysis of Smo loss of function clones. (A-C) In contrast to control clones (A, red), clones expressing a dominant negative Smo construct (SmoDN) (B, red) are largely eliminated from the Zfh1 positive (green) compartment containing the CySCs and their immediate progeny by 5d ACI. C) Quantification of the Zfh1 positive fraction amongst all clonal cells. Each data point represents one testis. Bar marks median, ***, p<0.001, Mann-Whitney U-test. (D-F) Clones (red) homozygous for the loss of function allele *rel*^*E20*^ (D) are retained in the Zfh1 positive (green) compartment by 5d ACI similar to control clones. Concomitant loss of rel (E) suppresses the elimination of clones (red) expressing SmoDN. (A,B,D,E) Cell outlines and hub marked by Armadillo (Arm) (blue), filled and open arrowheads mark Zfh1 positive and negative clonal cells, respectively, scale bars 10μm. (F) Quantification of the Zfh1 positive fractions per testis in control and experimental clones. At 5d ACI, loss of rel by itself has no effect on clone retention in the Zfh1 positive compartment, but does rescue the elimination of clones with SmoDN overexpression. n, number of testes; bar, median; small square, mean; box, 1st/3rd quartile; whiskers, last data point within 1.5x interquartile distance; x, outliers; ***, p<0.001, Kruskal-Wallis test with Tukey’s HSD; n.s., not significant.

### Suppression of loser elimination by inactivation of Rel does not depend on the specific trigger for competition

We next tested whether the rescue of stem cell state by inactivation of Rel was specific to loser cells induced by clonal loss of Hh niche signalling. For this, we first interfered with the essential Jak/Stat niche signal conveyed by Upd by clonally overexpressing Latran, an endogenous decoy receptor that resembles the active Upd receptor Domeless, but lacks the cytoplasmic domain required for signal transduction (Makki et al., 2010). Cells with Upd signalling impaired by Latran were largely eliminated through differentiation already by 3 days ACI (Fig. 3A,C). Analogous to the SmoDN experiment we next compared these cells to *rel* MARCM clones overexpressing Latran, and again observed a rescue of the number of cells retaining the stem cell marker Zfh1 under Latran overexpression when Rel was concomitantly inactivated (Fig. 3B,C).

**Figure 3:**
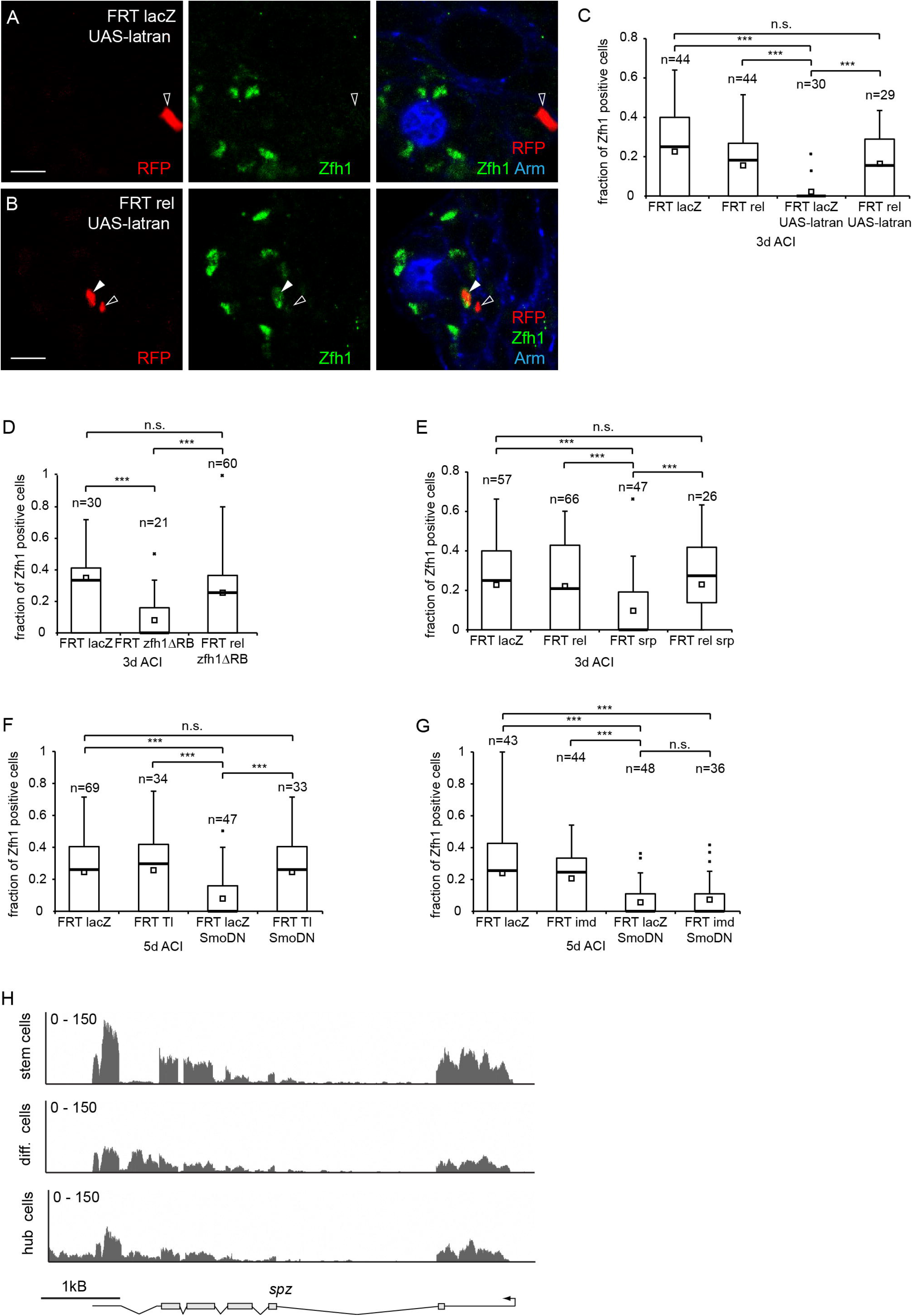
Rel dependent loser elimination is independent of the competition trigger and involves Tl but not Imd. (A-C) In contrast to control clones (red) (A), clones expressing the dominative negative Upd decoy receptor Latran (B) (red) are largely eliminated from the Zfh1 positive (green) stem cell compartment by 3d ACI. Cell outlines and hub marked by Armadillo (Arm) (blue), filled and open arrowheads mark Zfh1 positive and negative clonal cells, respectively, scale bars 10μm. (C-G) Boxplot quantifications of the Zfh1 positive fractions per testis in control and experimental clones. (C) Loss of Rel rescues the elimination of clones caused by Latran overexpression. (D) Loss of Rel is able to suppress the elimination of homozygous clones lacking the long Zfh1 isoform Zfh1-RB or the *Drosophila* GATA factor Srp (E) at 3d ACI. (F) Homozygosity for the Tl loss of function allele *Tl*^*RXA*^ has no effect on the fraction of clones retained in the Zfh1 positive compartment at 5d ACI, but rescues the elimination of clones overexpressing SmoDN. (G) Elimination of SmoDN overexpressing clones is not sensitive to inactivation of Imd. (H) RNAseq reveals expression of Spz in isolated CySCs (top track), differentiated CyCs (centre track), and hub cells (bottom track). Number of reads plotted against genome position. (C-G) Boxplots: n, number of testes; bar, median; small square, mean; box, 1st/3rd quartile; whiskers, last data point within 1.5x interquartile distance; x, outliers; ***, p<0.001, Kruskal-Wallis test with Tukey’s HSD; n.s., not significant.

To add another condition where elimination of clones is unambiguosly due to competition we turned to an isoform specific deletion of Zfh1, *zfh1*^*Δ*RB^. Both the short isoform Zfh1-RA and the longer variant Zfh1-RB that contains two extra N-terminal zinc fingers are expressed in the testis (Albert et al., 2018). However, while a deletion of Zfh1-RA is lethal, *zfh1*^*Δ*RB^ flies (Fig. S1C) are homo- or hemizygous viable and fertile. The Zfh1-RB isoform is therefore in principle dispensable for somatic stem cell function in the testis. Correspondingly, testes of flies carrying *zfh1*^*Δ*RB^ over a deficiency deleting the entire locus were morphologically normal and contained normal numbers of cyst stem cells marked by the remaining isoform Zfh1-RA (Fig. S1D). Nevertheless, both homozygous *zfh1*^*Δ*RA^ and *zfh1*^*Δ*RB^ clones were eliminated from the stem cell pool (Fig. S2B,C), in case of *zfh1*^*Δ*RB^ thus demonstrably due to competition. Concomitant clonal inactivation of Rel using a FRT82B *rel*^E20^ *zfh1*^*Δ*RB^ recombinant chromosome again enabled mutant clones to remain in the Zfh1 positive pool (Fig. 3D, Fig. S2D).

Similarly, while clones homozygous for the Srp loss of function allele *srp*^PZ01549^ are largely lost from the stem cell / early CyC pool by 3d ACI (Hof-Michel and Bökel, 2020), homozygous clones for an FRT *rel*^E20^ *srp*^PZ01549^ double mutant chromosome arm were retained in the Zfh1 positive compartment at a significantly higher fraction (Fig. 3E, Fig. S2E,F).

Thus, loss of Rel is able to suppress the elimination of loser clones from the stem cell compartment in response to a variety of different triggers used to induce loser fate.

### Cell competition can be suppressed by loss of Tl but not by inactivation of Imd

We next tested which signalling pathway was acting upstream of Rel in our somatic stem cell competition scenario. We mapped the molecular lesion in the genetically amorphic allele *tl*^RXA^ (Anderson et al., 1985) to a 4 aa deletion in an evolutionary conserved stretch at the end of the intracellular TIR domain (Fig. S3A) and recombined it onto an FRT82B chromosome for clonal analysis. We also generated a FRT42D chromosome carrying the Imd loss of function allele *imd*^10191^ (Pham et al., 2007), which abolishes signalling through the other branch of the innate immune response. As with Rel, clonal loss of Tl or Imd function had, by itself, no influence on the retention of the mutant cells in the Zfh1 positive stem cell / early CyC compartment (Fig.3F,G; Fig. S3B,D). Importantly, concomitant loss of Tl (Fig. 3F; Fig. S3C) but not Imd function (Fig. 3G; Fig. S3E) was able to suppress the competition induced loss of SmoDN overexpressing clones from the Zfh1 positive, stem cell containing pool. Thus, Rel function during competition in the testis does not depend on the canonical upstream pathway involving surface peptidoglycan receptors and the adaptor molecule Imd that operates in immune signalling. Rel activation thus appears to be controlled by a non-canonical, competitive signalling module downstream of Tl, resembling the situation in the wing disc (Alpar et al., 2018; Meyer et al., 2014). Both the competing stem cells and their somatic neighbours also transcribe the Tl ligand Spz (Fig. 3H). Again as in the larval wing disc (Alpar et al., 2018; Meyer et al., 2014), stem cell competition in the testis may therefore involve a local source of the ligand triggering the competitive signal.

### Rel is activated in cells adopting loser fate

Tl and Rel are required in loser cells for their elimination by differentiation from the Zfh1 positive stem cell pool, but expression of antimicrobial target genes indicating Rel activation was not observed near the testis tip under steady state, physiological conditions (Fig. 1C; Fig. S1B). We therefore asked whether Rel was instead activated in somatic stem cells only upon adopting loser fate. To test this hypothesis we generated a transcriptional sensor for Rel activity, placing a synthetic pomotor consisting of a tandem triplicated consensus Rel binding site upstream of a Hsp70 minimal promoter in front of a GFP-nls ORF (3xRelBS-GFPnls) (Fig. S4A).

We validated this reporter using a second transgenic construct comprised of a UAS promoter driving expression of a truncated Rel protein containing only the transcriptionally active, 68 kDa N-terminal fragment of Rel, but not the C-terminal, autoinhibitory domain (UAS-Rel68) (Fig. S4B). Driving UAS-Rel68 expression in the cyst cell lineage under tj-Gal4 tub-Gal80^ts^ control over night caused expression of GFPnls from the 3xRel promoter in both Zfh1 positive and negative cells (Fig. 4A,B). The same Rel68 expression also activated an established 3xRelBS>RFP_stop>luc construct (Chen et al., 2016) (Fig. S4C,D). Thus, our 3xRelBS-GFPnls reporter was in principle able to detect the presence of active Rel.

**Figure 4:**
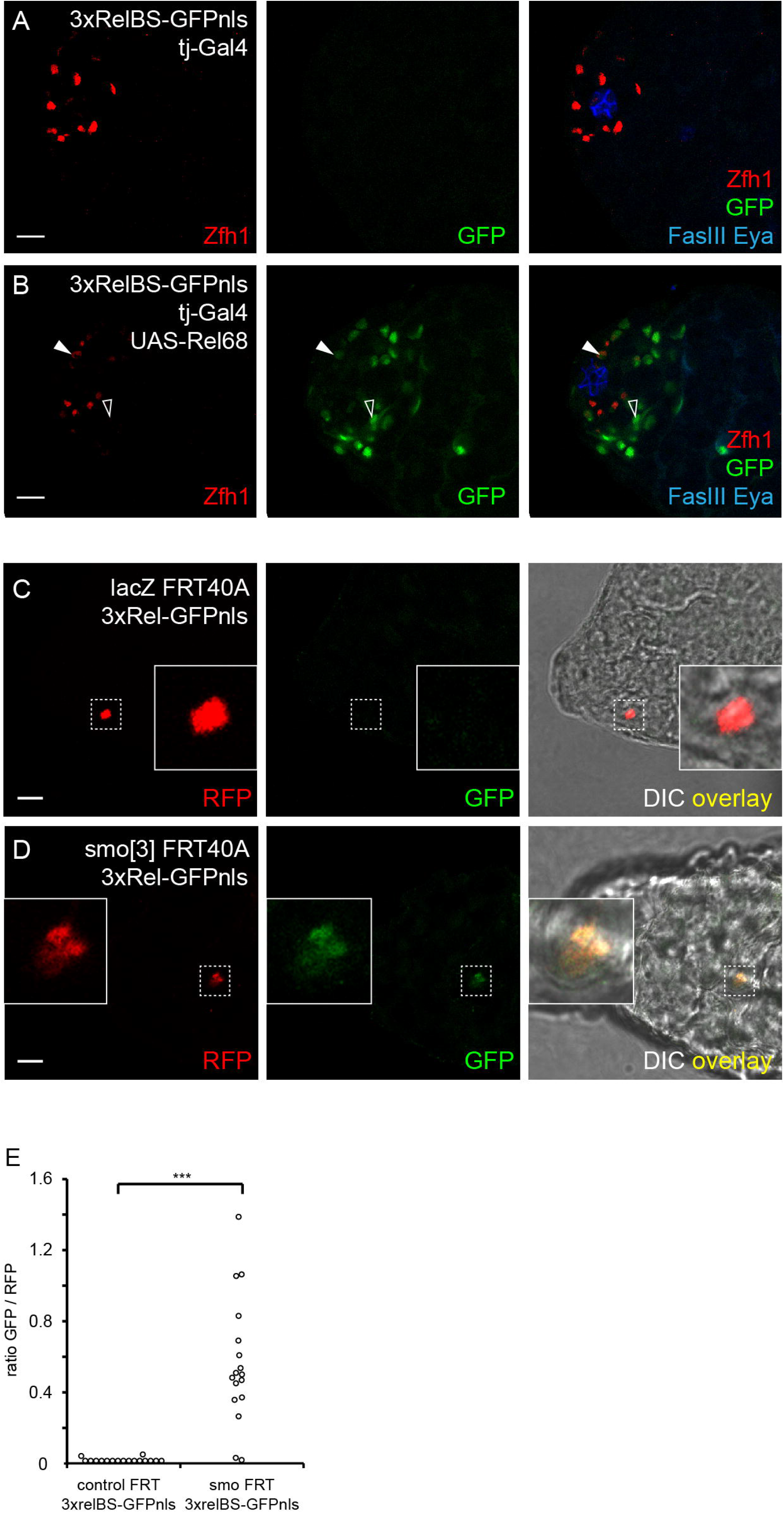
Rel is activated in loser cells during stem cell competition. (A,B) GFP expression (green) from a 3xRel binding site reporter is absent from driver only control testes (A) but is readily detected in both Zfh1 (red) positive (solid arrowheads) and negative (open arrowheads) cells after expressing UAS-Rel68 under tj-Gal4 control (B). Hub marked by FasIII and differentiated cells by Eya, blue. (C,D) In fixed, unstained testes, neutral MARCM clones (red) exhibit no 3xRel-GFP reporter activity (C), while reporter fluorescence is readily detectable in *smo* loser clones at 36h ACI (D). (E) Quantification of 3xRel-GFP reporter fluorescence normalized against the MARCM RFP signal in control and Smo clones. ***, p <0.001, Mann-Whitney U-test. (A-D) Scale bars, 10μm.

We next generated nlsRFP marked control and *smo*^3^ loser clones in a 3xRelBS-GFPnls background, and quantified GFP fluorescence in the clones by imaging the testes directly after dissection and fixing at 36h ACI, which is around the earliest timepoint when MARCM clones in the somatic lineage of the testis can be visualized. While the GFP signal in most control clones was indistinguishable from background, nuclei of *smo*^*3*^ clones showed significantly increased GFP fluorescence levels (Fig. 4C-E), indicating Rel activation specifically in loser cells.

### Rel activation triggers clone elimination independently from differentiation

The experiments described above established that in intact testes Rel activation occurs specifically in the losers of stem cell competition, and that Rel and Tl are required in these cells for their elimination and differentiation. To test whether Rel activation is also sufficient for executing loser fate we expressed the constitutively active Rel68 N-terminal fragment in control clones, as well as in *ptc*^*IIW*^ MARCM clones that constitutively activate Hh signalling, continue to express Zfh1 and proliferate also outside the niche, and act as winners by skewing neutral cell competition (Amoyel et al., 2013; Michel et al., 2012).

Neutral clones expressing activated Rel were retained within the Zfh1 positive pool containing the stem cells at significantly lower levels than control clones (Fig. 5A,B,G), demonstrating that Rel activation is sufficient for their elimination. In contrast, the fraction of Zfh1 positive cells did not differ significantly between *ptc*^*IIW*^ clones and *ptc*^*IIW*^ clones overexpressing activated Rel (Fig. 5C,D,G).

**Figure 5:**
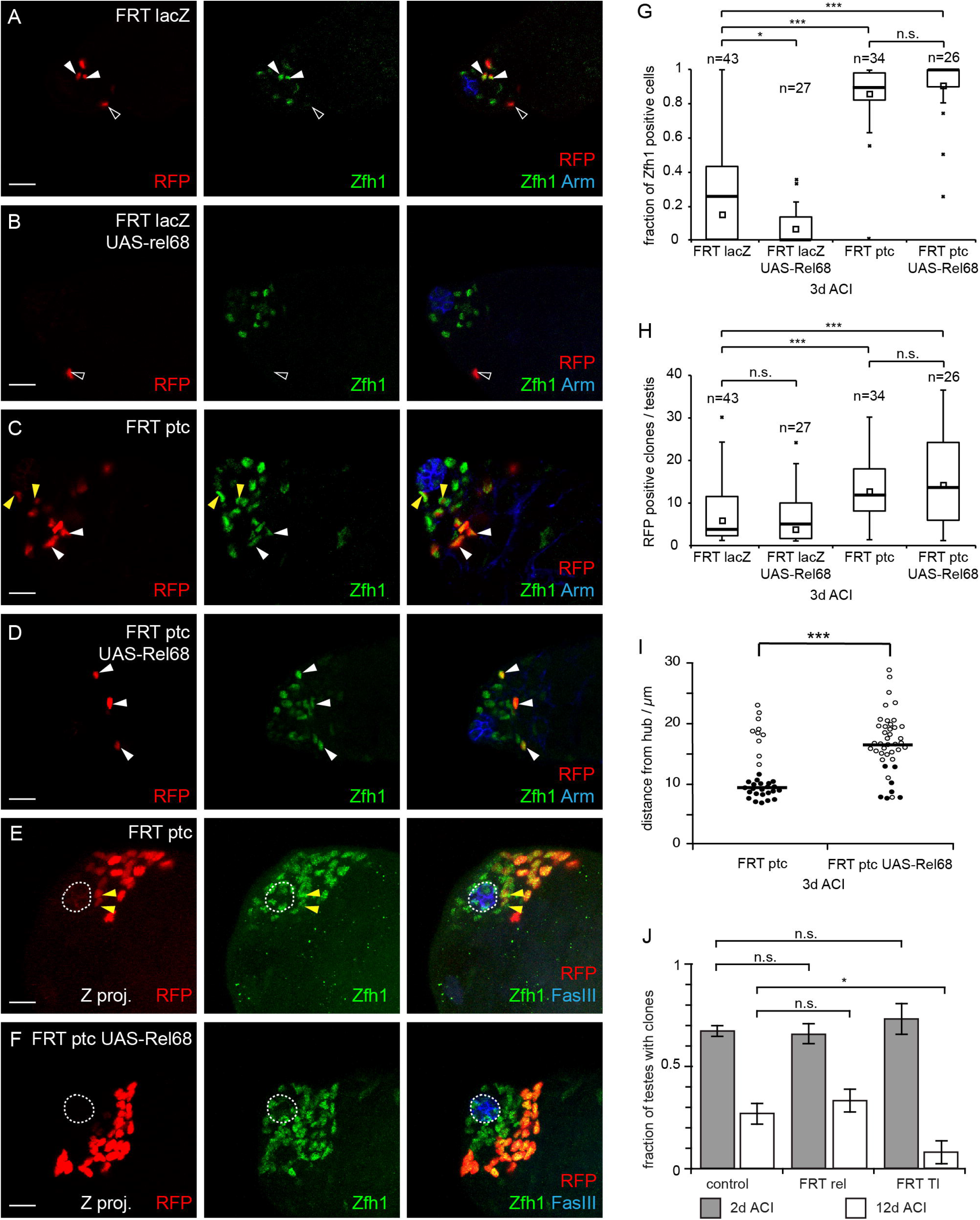
Rel activation is sufficient for the displacement of neutral and winner cells from the niche but Rel and Tl do not affect long term clone retention. (A-F) Control and *ptc* MARCM clones (red) with and without UAS-Rel68 overexpression. (A) In control testes, clonal cells are found within (solid arrowheads) and outside (hollow arrowheads) the Zfh1 positive (green) stem cell compartment. (B) Co-overexpression of Rel68 eliminates control MARCM clones from the stem cell pool. (C,D) Testes with *ptc* MARCM controls contain an elevated number of clonal cells, most of which remain Zfh1 positive (solid arrowheads) even outside the niche. Note the presence of Zfh1 positive, *ptc* mutant cells remaining in the immediate vicinity of the hub and embedded within non-clonal, Zfh1 positive cells (yellow arrowheads) in *ptc* clones not overexpressing Rel68 (C). (E,F) Z projections of FRT *ptc* (E) and FRT *ptc*; UAS-Rel68 (F) testes, Note the presence of clonal cells abutting the hub (yellow arrowheads) in (E), but the presence of a gap between the hub (dashed outline) and the innermost clones in (F). (A-F) Hub labelled with Arm, blue, scale bar 10 μm. (G,H) Quantification of the Zfh1 positive fraction of all clonal cells (G) and the absolute number of clonal cells per testis (H) in control and *ptc* MARCM clones with and without UAS-Rel68 overexpression. (I) Quantification of the distance from the innermost clonal cell to the centre of the hub. Each data point corresponds to one testis. Testes where the innermost clonal cell was embedded within the non-clonal stem cell population are indicated in solid black. (J) Fraction of testes containing RFP positive control, *rel*^*E20*^ or *Tl*^*RXA*^ at 2d and 12d ACI. Bars, mean ± SD, n=3 experiments. (G-I) bar, median. (G,H) n, number of testes; small square, mean; box, 1st/3rd quartile; whiskers, last data point within 1.5x interquartile distance; x, outliers. ***, p<0.001 and *, p<0.01, Kruskal-Wallis (G,H) and ANOVA (J) tests with Tukey’s HSD, Mann-Whitney U-test (I); n.s., not significant.

However, regular *ptc*^*IIW*^ clones are retained within the niche and, over time, outcompete and displace their wildtype neighbours from the immediate vicinity of the hub (Amoyel et al., 2013; Michel et al., 2012). Consistently, RFP labelled *ptc*^*IIW*^ clones were generally also present within the first tier of Zfh1 positive cells surrounding the hub (Fig. 5C). Concomitant expression of activated Rel reduced the fraction of testes in which the closest *ptc*^*IIW*^ clone was embedded within the endogenously Zfh1 positive cells. This displacement was even more striking in Z projections of the confocal stacks used for quantification (Fig. 5E,F), and was reflected by an increase in the distance between the innermost RFP marked cell and the centre of the hub (Fig. 5H). Importantly, both the number of homozygous *ptc*^*IIW*^ clonal cells per testis (Fig. 5I), and the Zfh1 positive fraction among these cells (Fig. 5G) remained elevated over control irrespective of the presence or absence of activated Rel. Thus, ectopic expression of activated Rel is sufficient to induce displacement of the *ptc*^*IIW*^ winner clones from the hub, but does not affect the retention of these cells in an otherwise stem cell like state.

### Loss of Rel or Tl does not promote long term clone retention

Stem cell turnover due to stochastic, neutral competition is a hallmark of many stem cell / niche systems, including the somatic lineage of the *Drosophila* testis (Amoyel et al., 2014). The involvement of Tl and Rel in the elimination of losers of cell competition in the narrow sense raised the possibility that the same cassette also affected the neutral competition operating in the background. Similar to manipulation of Hippo/Wts/Yki signalling (Albert et al., 2018; Amoyel et al., 2014), loss of Tl or Rel might bias the ongoing, neutral competition in favour of the mutant cells, resulting in a smaller fraction of testes stochastically losing their clones. We therefore assayed clone retention by quantifying the fraction of testes containing RFP marked clones at 2d and 12d ACI. At the early time point, we observed no significant differences between control, *rel*^*E20*^, and *Tl*^*RXA*^ clones, indicating comparable induction frequencies. As expected due to neutral competition alone, the fraction of testes containing control clones had dimished by 12d ACI. This drop was also present in the testes with *rel*^*E20*^ clones, and even exacerbated in the case of *Tl*^*RXA*^ (Fig. 5J). Loss of Tl or Rel therefore does not appear to benefit the mutant cells in the context of neutral competition.

## Discussion

In summary, we have demonstrated that activation of the NF-κB family transcription factor Rel occurs during somatic stem cell competition in the testis, and is necessary for elimination of the loser cells from the niche. Rel activation in the stem cells appears to be regulated by a non-canonical, Tl dependent competitive signalling module that may be activated through Spz ligand that is strongly expressed in the CySCs but also hub and differentiated somatic cells (Hof-Michel and Bökel, 2020). Such local ligand production and pathway activation would follow the paradigm of myc-dependent supercompetition in the wing disc epithelium that leads to expression of the pro-apoptotic protein Hid (Alpar et al., 2018; Meyer et al., 2014). However, in the testis there is no evidence for increased apoptosis of loser cells, which instead undergo differentiation (Michel et al., 2012).

Expression of activated Rel was also sufficient for the displacement of both neutral clones and “winner” cells lacking the inhibitory Hh receptor Ptc from the niche, even though the latter cells, due to the constitutive activation of the Hh niche signalling pathway, retain other stem cell characteristics, including increased clone size reflecting their proliferative state and expression of the stem cell marker Zfh1. The most parsimonious explanation for this may be that adhesion to the hub is the primary target of Rel activation, with differentiation of the loser cells a secondary consequence of their exit from the niche. Such a model would also be consistent with the observation that Robo2/Slit signalling influences the competitive ability of cyst stem cells by regulating DE-Cadherin mediated adhesion to the hub (Stine et al., 2014), which had previously been shown to be essential for stem cell maintenance (Voog et al., 2008). A link between stemness and adhesion under physiological conditions also exists in the germline, where miRNA induced downregulation of N-Cadherin controls the detachment from the niche that is essential for stem cell turnover and allows GSCs to differentiate into gonialblasts (Epstein et al., 2017).

The competitive ability of somatic stem cell in the testis also depends on MAPK signalling (Amoyel et al., 2016; Singh et al., 2016), and it will be interesting to dissect the relationship between the various pathways, in particular since artificially increased adhesion to the hub mediated by integrin overexpression is able to suppress MAPK controlled expulsion of loser cells from the niche (Singh et al., 2016).

Rel is involved in the elimination of stem cells whose fitness has been compromised by interfering with a variety of different pathways or transcription factors, and thus appears to occupy a central role in integrating and regulating somatic stem cell competition in the testis. Triggers for competition we have identified include reduced signalling activity through the Hh and Upd - Jak/Stat niche signalling pathways, loss of the Drosophila GATA factor Srp that we recently identified as a stem cell factor in the cyst cell lineage (Hof-Michel and Bökel, 2020), and loss of the long isoform of the stem cell marker Zfh1 (Zfh1-RB). Since the complete absence of this isoform in homozygous *zfh1*^*ΔRB*^ animals has no effect on viabilty and fertility, the forced differentiation of individual stem cells lacking Zfh1-RB is unambiguosly due to competition. In an interesting parallel, in muscle satellite cells in *Drosophila* the long RB isoform is physiologically downregulated by miRNAs to induce differentiation, while the RA isoform lacking the seed site is refractory and Zfh1-RA expressing cells are thus retained as stem cells (Boukhatmi and Bray, 2018).

Zfh1 and Rel also interact during the innate immune response, where Zfh1 appears to act at the same level or downstream of Rel to downregulate Imd/Rel target genes such as CecB (Myllymaki and Ramet, 2013). In contrast, in the testis expression of CecB and other antimicrobial peptides, presumably triggered in response to bacterial exposure during cell isolation, is restricted to the Zfh1 positive pool of stem cells and their earliest progeny that also exhibit increased Rel levels (Hof-Michel and Bökel, 2020). It will be interesting to disentangle the relation between Zfh1 and Rel in the testis using approaches not suffering from crosstalk with active immune responses.

In conclusion, we have shown that activation of Rel downstream of a Tl dependent competitive signalling module is, during stem cell competition in the somatic lineage of the *Drosophila* testis, necessary and sufficient for the elimination of loser cells from the stem cell pool.

The ability of Rel to force loser cells into differentiation, irrespective of the nature of the fitness cues triggering loser fate, could explain why the loss of any one niche signal typically causes overall differentiation, rather than affecting only specific subsets of stem cell behaviours. This all-or-nothing response had been taken as evidence for a binary decision between stemness and differentiation. Adding competition to the list of hallmark stem cell properties instead lends further support to models that consider stemness to be a compound physiological state, comprised of independently regulated activities that contribute to self renewal and the generation of differentiating progeny (Clevers and Watt, 2018; Post and Clevers, 2019).

## Materials and Methods

### Fly stocks and transgenic constructs

w; arm-lacZ FRT40A (BL-7371), w;FRT42D arm-lacZ (BL-7372), w;;UAS-RedStinger (BL-8547), w;;FRT82B w^+^90E (BL-2050), wg^Sp^/CyO;3xRelBS-FRT-mCherry.stop-FRT-Luc (BL-67137), pBAC-RelGFP (BL-43956), and w;;Df(3R)exel9020/TM6B, Tb (BL-7917) were obtained from the Bloomington Drosophila Stock Center and have been described. The w hs-FLP C587-Gal4 UAS-RedStinger (Michel et al., 2012) and UAS-smoIPSA (Kupinski et al., 2013) (here referred to as UAS-SmoDN) chromosomes have been described before.

The following stocks were graciously provided by our colleagues: P{ry[+t7.2]=PZ} srp^01549^/ TM3,ry Sb Ser (srp^PZ^, Ingolf Reim, FAU Erlangen), w;;rel^E20^/TM6C, w;;attp2 Mtk-GFP/TM6C and w;;attP2 Drs-GFP (Marc Dionne, Imperial College London), UAS-latran (Michel Crozatier, CBI Toulouse), and ru h th e Tl^RXA^/TM6C (Bruno Lemaitre, EPF Lausanne).

To generate zfh1ΔRA and zfh1ΔRB we cloned two guide RNAs flanking the respective start codons into pU6-BbsI-chiRNA (Addgene #45946). For zfh1ΔRA, the two plasmids were directly co-injected into y w vas-Cas9 flies, and candidate mutants recovered by screening for lethality over Df(3R)Exel9020. For zfh1ΔRB we tandem inserted the U6 expression cassettes driving the two gRNAs into pCaSPer4. Following standard P transgenesis one of the resulting lines was crossed to y w vas-Cas9, and the resulting progeny screened by PCR and sequencing for the presence of the desired deletion.

For the pattB 3xRelBS-GFPnls construct we combined a tandem triplicated Rel binding site and Hsp70 minimal promoter with the GFPnls ORF taken from pSav-GFPnls (Albert et al., 2018) and the 3’UTR and polyadenylation site from a pUAST vector modified to also contain an attB cassette. Inserting the resulting construct into the attP2 landing site yielded the w;;3xRelBS-GFPnls reporter line. w;;UAS-rel68 was generated by placing a PCR product encoding the first 545 aa of the Rel ORF followed by a stop codon into pUAST attB and inserting the resulting construct into the attP2 landing site.

MARCM clones (Lee and Luo, 1999) were generated by crossing FRT males, where required also carrying additional UAS or reporter constructs, to w hs-FLP C587-Gal4 UAS-RedStinger virgins carrying the matching tub-Gal80 FRT chromosome and heat shocking adult males for 30min h at 37°C. Clones were generated using the following FRT chromosomes: arm-lacZ FRT40A, smo^3^ FRT40A, FRT42D arm-lacZ, FRT42D ptc^IIW^, FRT42D imd^10191^, FRT82 w^+^90E, FRT82 rel^E20^, FRT82 Tl^RXA^, FRT82 srp^PZ01549^, FRT82 rel^E20^ srp^PZ01549^, FRT82 zfh1^ΔRB^, FRT82 rel^E20^ zfh1^ΔRB^ that were generated, where required, by standard meiotic recombination between mutations and FRT sites followed by PCR genotyping.

Full sequences for all constructs are available upon request.

### Antibodies and immunohistochemistry

Testes were stained as described (Albert 2018). The following antisera were used: rat-anti-DECadherin (DSHB, DCAD2) 1:100, mouse-anti-FasIII (DSHB, 7G10) 1:250, mouse-anti-Armadillo (DSHB, N2 7A1) 1:100, rabbit-anti-Zfh1 (Ruth Lehmann) 1:4000. Goat secondary antisera labelled with Alexa-488, −568, or −633 (Invitrogen) were used 1:500.

### Imaging and image analysis

Images were acquired using a Leica SP5 confocal microscope with a HCX PL APO 40x / NA1.25 oil immersion objective. Unless stated otherwise, images are single optical slices. Image quantifications were performed using Fiji (Schindelin et al., 2012). Differences between samples were tested for significance as appropriate with Mann-Whitney U, ANOVA or Kruskal-Wallis tests followed by Tukey’s HSD, using the PMCMRplus R package (Pohlert, 2018). Images were prepared for publication using Adobe Photoshop and Illustrator.

### RNAseq

The transcriptome analysis of somatic cell populations in the fly testis has been described in detail in a separate manuscript (Hof-Michel and Bökel, 2020). In brief, cells were labelled *in vivo* by expressing the RedStinger nuclear RFP under appropriate Gal4 control. Following dissection of the testes, labelled cells were mechanically and enzymatically isolated and then sorted by FACS at the Flow Cytometry Core Facility at Philipps-University Marburg. mRNA was isolated from the samples, reverse transcribed, and amplified by the Deep Sequencing Core Facility of the Center for Molecular and Cellular Biolengineering at the Technical University Dresden and then sequenced using an Illumina NextSeq500 sequencer. For analysis of the raw data we used the A.I.R. RNAseq web based analysis package (Sequentia Biotech, Barcelona). The underlying algorithms and software packages are described in (Vara et al., 2019). For GO enrichment analyses we used the web based shinyGO platform (Ge et al., 2019).

## Supporting information

Table S1 Immune genes upregulated in CySCs vs. diff CyCs

## Author contributions

SH-M established the cell purification protocol and performed initial fly experiments, imaging, and analyses of imaging data, SH performed molecular cloning and fly genetics experiments, LC performed immunostaining and fly genetics experiments. CB performed fly and imaging experiments, was responsible for imaging and data analysis, devised the project, and wrote the manuscript.

## Acknowledgements

We would like to thank Marc Dionne, Marc Amoyel, Michel Crozatier, Ruth Lehmann, Bruno Lemaitre, and the JEDI community for fly stocks, reagents, and advice, Erika Bach for critical comments on the manuscript, Isolde Kranz for fly food and coffee, and Olga Puretskaja and Olga Sidorova for their work on the Zfh1 isoform specific deletions. The project was supported by DFG grant BO 3270/3-1 to CB.

