## Supplementary Information and Figures for "Innate immune signalling drives loser cell elimination during stem cell competition in the *Drosophila* testis"

**Supplementary Table S1**

**Supplementary Figures S1 to S4**

**Supplementary Table S1: Enrichment of immunity associated genes in CySCs vs. CyCs.** Genes significantly overexpressed in Cyst Stem Cells relative to their differentiated progeny (Hof-Michel and Bökel, 2020) are enriched for association with the GO term GO:0002376 "Immune System Process" (GO:0002376, 139/500 genes, FDR = 4.3 E-11). Genes directly involved in Tl or Imd signalling are highlighted in yellow.

**Figure S1: Drs expression and isoform specific Zfh1 deletions.** (A) RNAseq reveals expression of the antimicrobial peptide Drs in isolated CySCs (top track) but not differentiated CyCs (bottom track). (B) In testes fixed immediately after dissection a Drs-GFP reporter (green) is not expressed in the vicinity of the hub (marked by DE-Cad, blue) or the Zfh1 positive (red) pool comprised of CySCs and their immediate, differentiated cyst cells progeny. (C) Isoform specific deletions of Zfh1-RA and Zfh1-RB. Both deletions were generated by Crispr/Cas9. *zfh1*$^{\Delta RA}$ was recovered by injecting two gRNAs flanking the start codon into y w vasa-Cas9 and testing candidate chromosomes for lethality over Df(3R)exel9020 before PCR screening. *zfh1*$^{\Delta RB}$ is viable and was recovered by PCR screening progeny from crossing a dual gRNA transgenic chromosome to the Cas9 source. (D) *zfh1*$^{\Delta RB}$ is viable and fertile over Df(3R)exel9020. Testes of such males are morphologically normal, and Zfh1

labelled with DE-Cadherin (blue), scale bars 10 µm.

**Figure S2: SmoDN validation and Rel dependent elimination of *zfh1*$^{\Delta RB}$ and *srp*$^{PZ}$ clones** (A) The number of Zfh1 positive cells per testis is only marginally affected when SmoDN is expressed in all cells of the cyst cell lineage under tj-Gal4 Gal80$^{ts}$ control for 3d. (B-D) (C-E) Both FRT *zfh1*$^{\Delta RA}$ (B) and FRT *zfh1*$^{\Delta RB}$ (C) clones are excluded from the Zfh1 positive compartment at 3d ACI, while FRT *rel zfh1*$^{\Delta RB}$ double mutant clones (D) are retained. (E,F) FRT *srp*$^{PZ}$ clones (E) are excluded from the Zfh1 positive compartment at 3d ACI, while FRT *rel srp*$^{PZ}$ double mutant clones (F) are retained. (A) n, number of testes; bar, median; small square, mean; box 1st/3rd quartile; whiskers, last data point within 1.5x interquartile distance; x, outliers; *, p<0.01, Kruskal-Wallis test with Tukey's HSD; n.s., not significant. (B-F) Filled and open arrowheads mark Zfh1 positive and negative clonal cells, respectively, scale bars 10µm. Hub marked in blue by FasIII (C,D) or Arm (E,F).

**Figure S3: Inactivation of Tl but not Imd suppresses loser cell elimination.** (A) Structure of the Tl transmembrane receptor protein. The loss of function mutation $Tl^{RXA}$ is an in frame deletion of 4 aa near the end of the intracellular TIR domain that is essential for signal transduction. The affected region is conserved in evolution in Tl proteins from Dipterans and Hymenopterans to hemimetabolous insects, *Drosophila* Tl-related proteins, Crustacean Tl proteins, and human cytokine receptors. (B-E) MARCM clonal analysis using RFP labeled clones (red) in the testis tip. Hub labelled with FasIII or Arm, blue, stem cells with Zfh1, green, scale bars 10µm. Open arrowheads indicate Zfh1 negative and closed arrowheads Zfh1 positive clonal cells. (B,C) Clonal homozygosity for FRT $Tl^{RXA}$ does not affect retention of clones in the Zfh1 positive stem cell compartment (B) and is able to rescue the exclusion of clones expressing SmoDN (C). (D,E) Clonal homozygosity for FRT $imd^{10191}$ does not affect retention of clones in the Zfh1 positive stem cell compartment either (D), but fails to rescue the elimination of clones expressing SmoDN (C).

**Figure S4: Validation of the 3xRelBS-GFPnls and UAS-Rel68 constructs.** (A) The 3xRelBS-GFPnls construct was generated by placing three consensus Rel binding sites (5'GGGAAACCCCCATTTGC3') in tandem upstream of a hsp70 minimal promoter and a GFPnls ORF and a mini white transgenesis marker. A downstream attB site was used to target the construct to the attP2 landing site. (B) Structure of the Rel NF-κB transcription factor. The N-terminal 545aa constitute the active, 68 kDa transcription factor (Rel68) with its IPT (Ig-like, Plexin, Transcription factor) domain, while the autoinhibitory C-terminal half possesses Ankyrin repeats and retains the uncleaved protein in the cytoplasm. The UAS-rel68 construct was generated by cloning a genomic fragment encoding the first 545aa of Rel and containing one intron into an pUAST attB. (C,D) A published 3xRelBS-FRT-RFP_Stop-FRT-luc construct generated no reporter fluorescence (red) in a tj-Gal4 driver only control (C), but yielded a signal in the vicinity of the hub (FasIII,blue) in the presence of UAS-Rel68 (D).

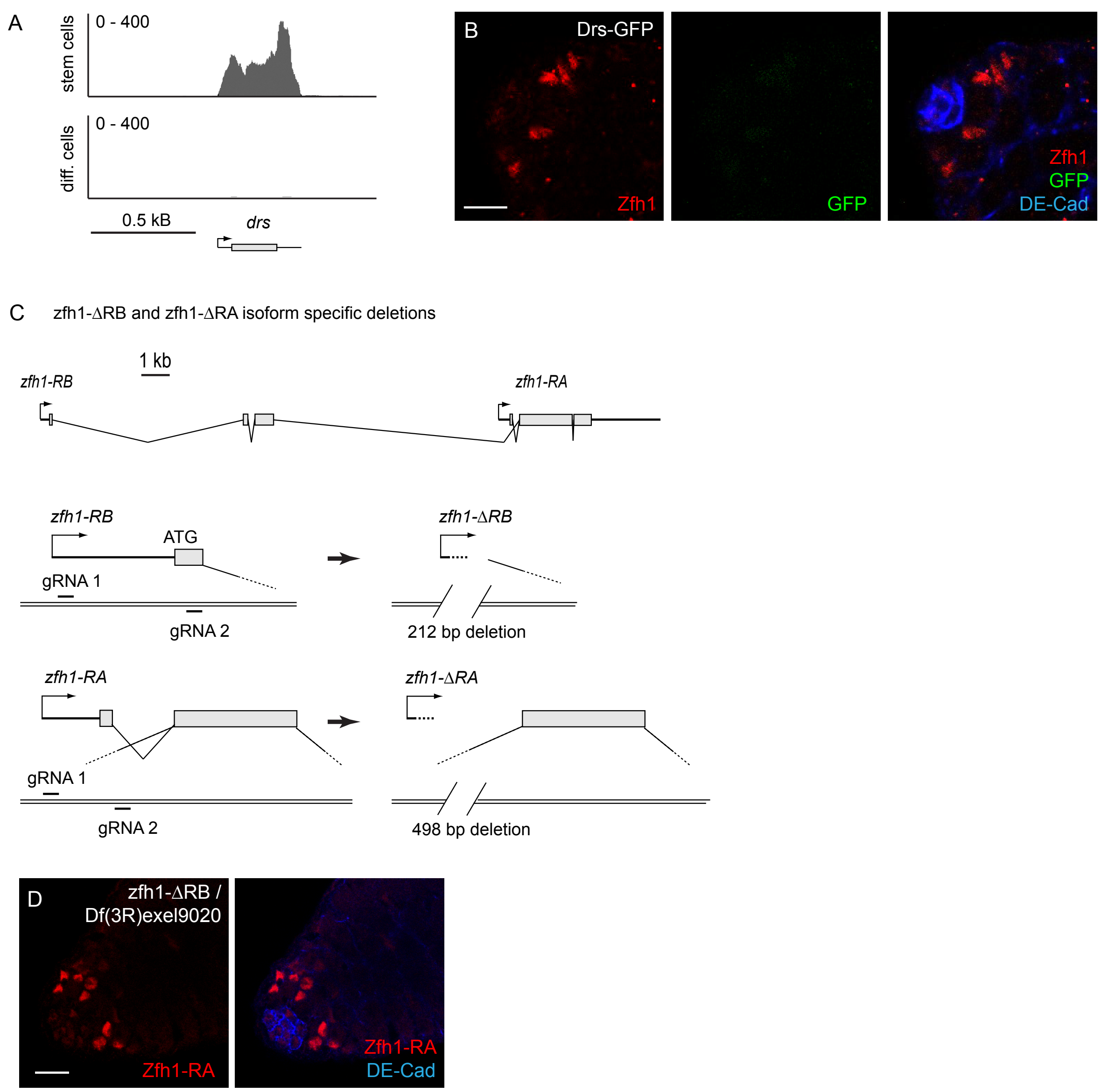
A
stem cells
0 - 400
diff. cells
0 - 400
0.5 kB
drs
B
Drs-GFP
Zfh1
GFP
Zfh1
GFP
DE-Cad
C
zfh1-ΔRB and zfh1-ΔRA isoform specific deletions
1 kb
zfh1-RB
zfh1-RA
zfh1-RB
ATG
gRNA 1
gRNA 2
zfh1-ΔRB
212 bp deletion
zfh1-RA
gRNA 1
gRNA 2
zfh1-ΔRA
498 bp deletion
D
zfh1-ΔRB /
Df(3R)exel9020
Zfh1-RA
Zfh1-RA
DE-Cad

**Fig. S1**

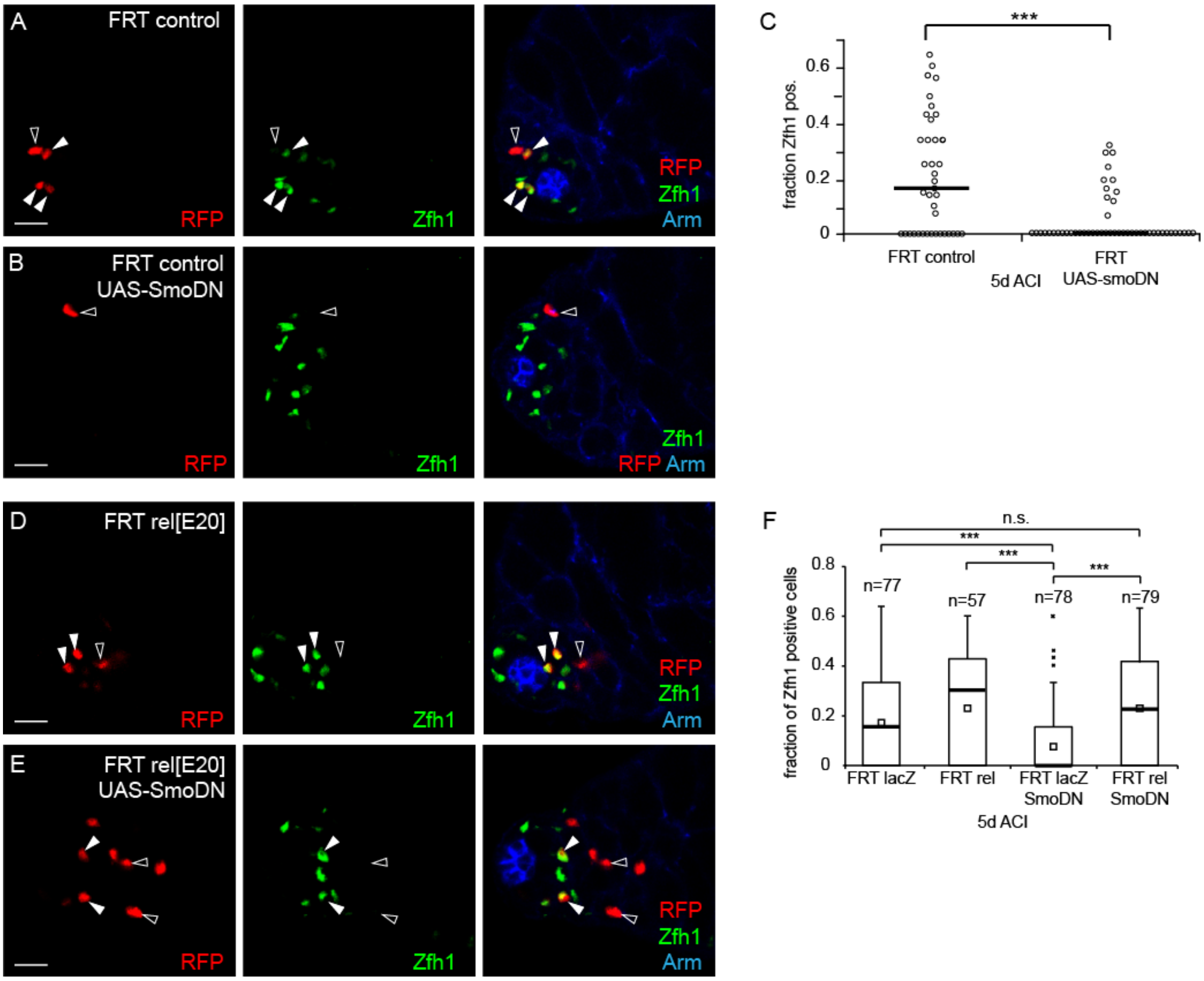

A
FRT control
RFP
Zfh1
RFP
Zfh1
Arm
B
FRT control
UAS-SmoDN
RFP
Zfh1
Zfh1
RFP Arm
C
***
fraction Zfh1 pos.
0.6
0.4
0.2
0
FRT control
5d ACI
FRT
UAS-smoDN
D
FRT rel[E20]
RFP
Zfh1
RFP
Zfh1
Arm
E
FRT rel[E20]
UAS-SmoDN
RFP
Zfh1
RFP
Zfh1
Arm
F
n.s.
***
***
***
fraction of Zfh1 positive cells
0.8
0.6
0.4
0.2
0
n=77
n=57
n=78
n=79
FRT lacZ
FRT rel
FRT lacZ
SmoDN
FRT rel
SmoDN
5d ACI


**Fig. S2**

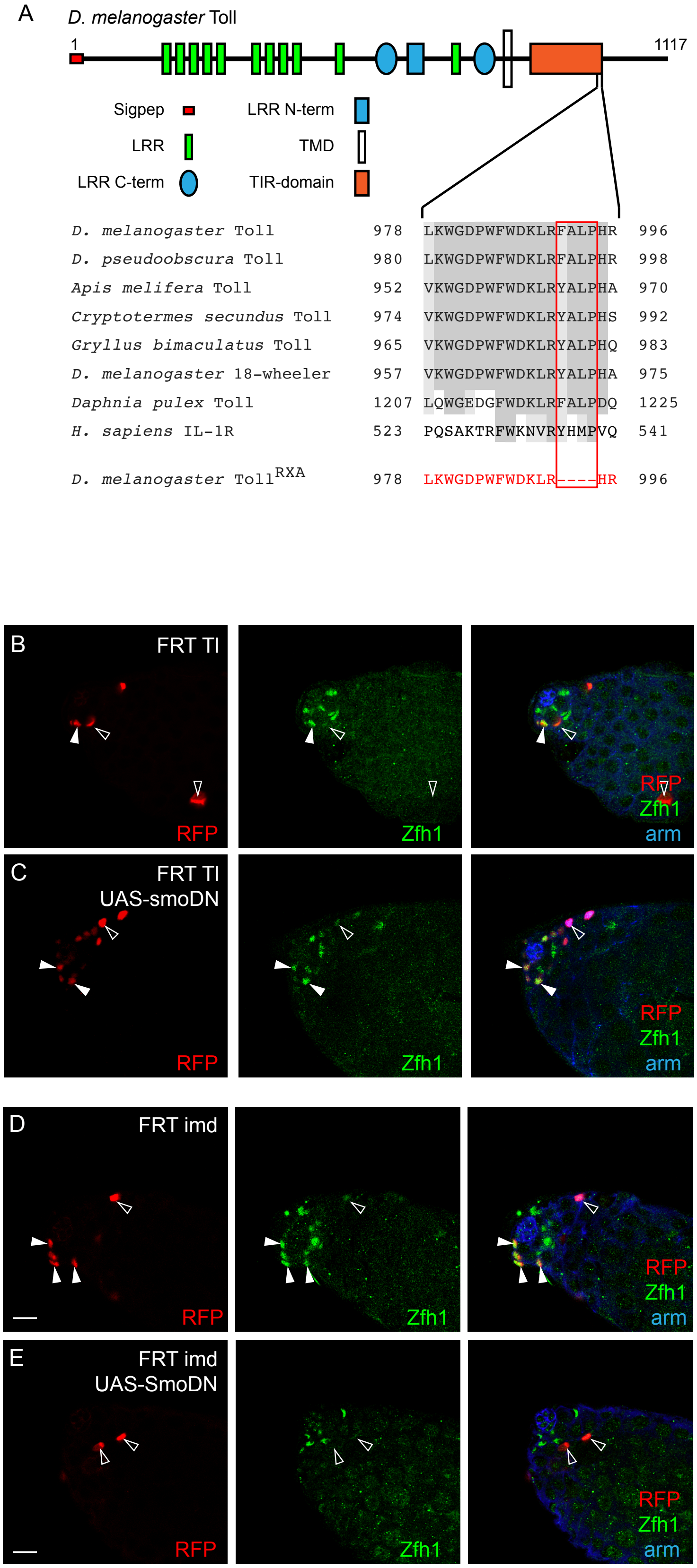

A
D. melanogaster Toll
1
1117
Sigpep
LRR N-term
LRR
TMD
LRR C-term
TIR-domain
D. melanogaster Toll 978 LKWGDPWFWDKLRFALPHR 996
D. pseudoobscura Toll 980 LKWGDPWFWDKLRFALPHR 998
Apis melifera Toll 952 VKWGDPWFWDKLRYALPHA 970
Cryptotermes secundus Toll 974 VKWGDPWFWDKLRYALPHS 992
Gryllus bimaculatus Toll 965 VKWGDPWFWDKLRYALPHQ 983
D. melanogaster 18-wheeler 957 VKWGDPWFWDKLRYALPHA 975
Daphnia pulex Toll 1207 LQWGEDGFWDKLRFALPDQ 1225
H. sapiens IL-1R 523 PQSAKTRFWKNVRYHMPVQ 541
D. melanogaster Toll^RXA 978 LKWGDPWFWDKLR----HR 996
B
FRT Tl
RFP
Zfh1
RFP
Zfh1
arm
C
FRT Tl
UAS-smoDN
RFP
Zfh1
RFP
Zfh1
arm
D
FRT imd
RFP
Zfh1
RFP
Zfh1
arm
E
FRT imd
UAS-SmoDN
RFP
Zfh1
RFP
Zfh1
arm


**Fig. S3**

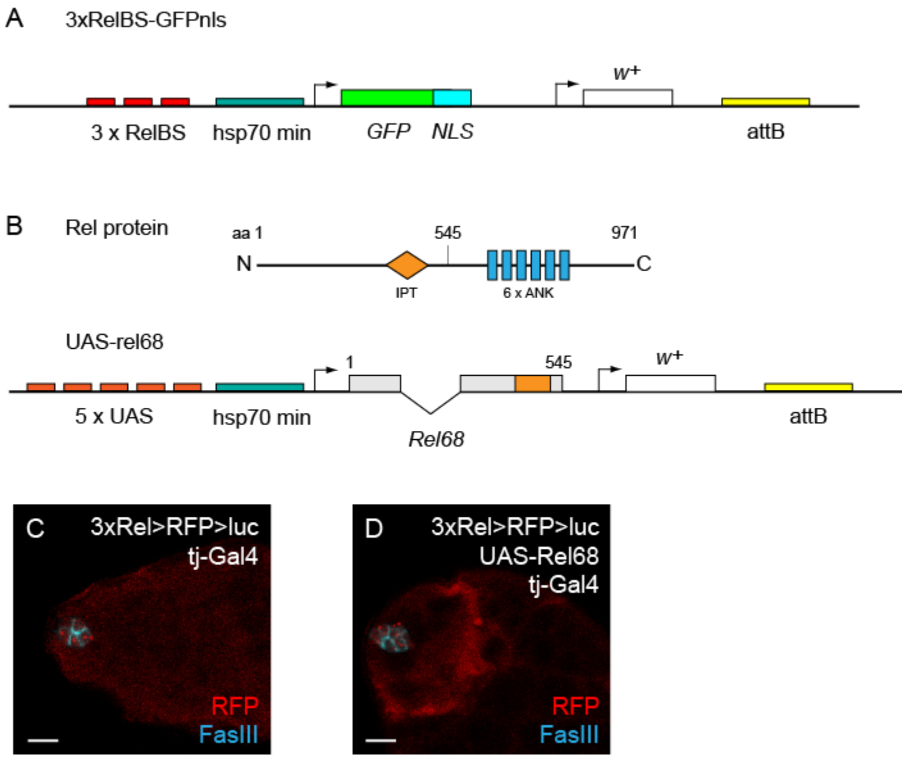
A
3xRelBS-GFPnls
w+
3 x RelBS
hsp70 min
GFP
NLS
attB
B
Rel protein
aa 1
545
971
N
C
IPT
6 x ANK
UAS-rel68
1
545
w+
5 x UAS
hsp70 min
Rel68
attB
C
3xRel>RFP>luc
tj-Gal4
RFP
FasIII
D
3xRel>RFP>luc
UAS-Rel68
tj-Gal4
RFP
FasIII

**Fig. S4**
